# BDNF receptor balance gates AgRP-to-PVH fiber density and stimulation-evoked feeding across caloric states

**DOI:** 10.1101/2025.09.12.675848

**Authors:** O. Yipkin Calhan, Sara K Michel-Le, Elizabeth N. Godschall, Shishir S Sriramoju, Margaret Nunnaly, Sophia Ogilvie, Kaleigh I. West, Orien Dong-Ang Li, Leann Vo, Taha Bugra Gungul, Prathik Reddy Nalamalapu, Marcos Torales Montes, Lloyd Choi, John N Campbell, Ali D. Güler, Christopher D. Deppmann

## Abstract

Agouti-related peptide (AgRP) neurons show opposite pairings of anatomy and output across caloric states. In mice, 24-h caloric restriction (CR) increases AgRP-to-paraventricular hypothalamus (PVH) fiber density without increasing stimulation-evoked feeding, whereas 5-day high-fat diet (HFD) reduces fiber density yet increases stimulation-evoked feeding, including during PVH terminal stimulation. PVH Bdnf rises acutely under both challenges and remains elevated with sustained HFD. Receptor expression shifts with state: in AgRP neurons, CR increases TrkB transcripts, whereas HFD increases p75NTR transcripts. AgRP-specific deletions define directional roles: TrkB loss reduces AgRP-to-PVH fiber density and yields larger stimulation-evoked feeding, whereas p75NTR loss increases fiber density and limits responses. Thus, a TrkB/p75NTR balance in AgRP neurons provides rheostat-like control over the relationship between circuit structure and stimulation-evoked feeding across caloric states, identifying TrkB and p75NTR as tractable molecular handles for probing state-dependent control of AgRP circuit output.

**Highlights:**

- CR increases AgRP→PVH fibers without boosting evoked feeding
- HFD retracts AgRP→PVH fibers and **amplifies** evoked feeding
- PVH **Bdnf** rises acutely in response to CR or HFD; elevation persists only with HFD
- AgRP **TrkB**↑ with CR, **p75NTR**↑ with HFD; AgRP specific knockouts of these receptors define floor and ceiling PVH axon innervation/evoked feeding

## Introduction

Maintaining energy balance requires the brain to integrate diverse metabolic cues and flexibly adjust feeding behavior in response to changing physiological demands [1]. The hypothalamus plays a central role in this regulation, acting as a key hub for sensing hormonal and nutrient-derived signals such as leptin, insulin, ghrelin, and glucose [2,3]. These signals are relayed to distinct hypothalamic neuron populations that coordinate food intake, energy expenditure, and overall homeostasis [4]. Disruption of this regulatory network contributes to the development of obesity and related metabolic disorders [5,6], making it essential to understand the mechanisms that allow hypothalamic circuits to adapt to sustained changes in caloric state.

Among these hypothalamic populations, agouti-related peptide (AgRP)-expressing neurons in the arcuate nucleus (ARC) are central to driving hunger and promoting food intake, especially during energy deficit [7] [8–10]. AgRP neurons are classically described as acute metabolic sensors, rapidly adjusting their firing activity in response to energy need and availability [11–15]. In contrast, far less is understood about how these neurons undergo longer-term structural and functional adaptations to sustained caloric challenges, such as during chronic caloric restriction (CR) or high-fat diet (HFD) exposure.

Although the underlying mechanisms remain obscure, AgRP neurons clearly possess the capacity for plasticity, undergoing rapid structural and functional remodeling in response to energy state. For example, fasting promotes dendritic spinogenesis and strengthens excitatory synaptic inputs onto AgRP neurons, enhancing their activity and driving feeding behavior [16]. These neurons also remodel their axonal projections to key downstream targets, such as the PVH, in response to energy deficits, a process regulated by ghrelin and other circulating factors [17]. In contrast, HFD feeding, both acute and chronic, elevates AgRP neuron activity, even before significant weight gain or leptin resistance, indicating that dietary composition can directly alter their excitability [18]. These adaptations are essential for maintaining energy balance, but sustained or maladaptive remodeling may reinforce persistent hunger and contribute to obesity-related disorders.

Neurotrophic signaling, particularly via brain-derived neurotrophic factor (BDNF), is an attractive candidate mechanism for mediating this plasticity. BDNF signaling is widely recognized for its role in activity-dependent neuronal remodeling, synaptic refinement, and circuit plasticity throughout the nervous system [19] [20]. In the hypothalamus, BDNF signaling critically regulates energy balance [21–28], and impaired BDNF function has consistently been associated with obesity in both animal models and humans [29,30]. BDNF acts primarily through two distinct receptors: tropomyosin receptor kinase B (TrkB), typically involved in promoting synaptic stabilization and neuronal survival [31], and the p75 neurotrophin receptor (p75NTR), known to mediate diverse cellular responses, including axonal pruning, synapse elimination, and functional modulation [32,33]

Our previous work has shown that p75NTR in AgRP neurons plays a critical role in anticipatory feeding behavior, suggesting that neurotrophin signaling contributes to both acute and predictive aspects of hunger [22,28]. However, whether BDNF receptor signaling enables AgRP neurons to flexibly remodel across different caloric states remains unclear. To address this, we examined how short-term caloric restriction and high-fat diet feeding alter AgRP→PVH connectivity and feeding behavior, whether BDNF expression in the PVH responds to these challenges, and how receptor-specific knockouts of TrkB or p75NTR affect AgRP neuron plasticity. These studies identify distinct roles for BDNF receptors in coordinating structural and functional adaptations of hypothalamic circuits under dynamic metabolic conditions.

## Results

### AgRP Neuron Projections to PVH Are Rapidly Strengthened by Caloric Restriction

Previous work has shown that acute fasting can induce rapid remodeling of AgRP neuron projections to the paraventricular hypothalamus (PVH), a key downstream node in the hunger circuit [17]. To determine whether similar remodeling occurs under calorie deficiency, we examined both anatomical and functional adaptations of AgRP neurons following 1- and 5-day caloric restriction (CR).

AgRP-IRES-Cre::tdTomato mice were used to visualize AgRP projections to the PVH under standard diet (SD), 1-day CR, and 5-day CR conditions (Figure 1A). The PVH was anatomically defined based on stereotaxic coordinates and cytoarchitectural landmarks in the Paxinos and Franklin mouse brain atlas. To capture region-specific changes along the rostrocaudal axis, we quantified AgRP fiber density and fluorescence intensity across the anterior, middle, and posterior PVH in 100-μm intervals. Following 1-day CR, a significant increase in AgRP innervation was observed specifically in the mid-PVH (approx. AP −0.70 to −0.94 mm from bregma), while no significant changes were detected in the anterior or posterior subregions (p < 0.05; Figure 1B–C). Although a similar trend was observed after 5-day CR, the increase did not reach statistical significance Given that 1-day CR increased AgRP→PVH innervation, we next asked whether this structural remodeling translates into functional changes in feeding behavior. To test this, we employed in vivo optogenetic activation of AgRP neurons, which allows for precise, temporally controlled stimulation and provides a standardized, causal measure of AgRP neuron output independent of upstream inputs or internal metabolic state. pAAV1-EF1a-DIO-hChR2(H134R)-EYFP-WPRE-HGHpA or control virus pAAV1-EF1a-DIO-EYFP was injected into the arcuate nucleus of AgRP-IRES-Cre mice, and an optical fiber was implanted 0.2 mm above the ARC (Figure 1D). Following three weeks of recovery, mice underwent either 1-day or 5-day CR (Figure 1E). To evaluate how CR influences the food-promoting effect of AgRP neurons, we measured stimulation-evoked feeding across a range of frequencies both before and after CR. This dose–response design provided a baseline curve for each animal, allowing us to directly assess whether CR potentiated or depressed the feeding response to AgRP activation.

**Figure 1.**
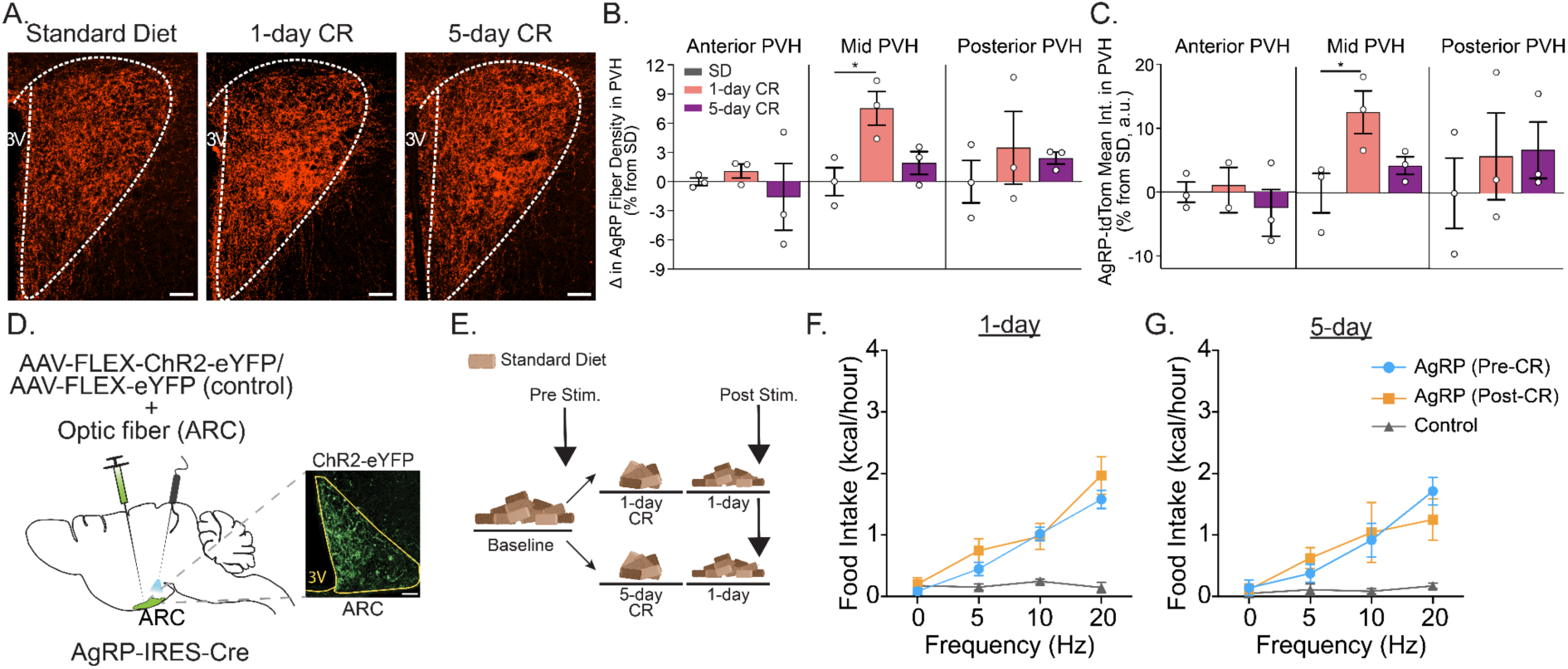
Caloric restriction increases AgRP→PVH innervation but does not potentiate optogenetically evoked feeding. **(A)** Representative images of AgRP-tdTomato-labeled fibers in the paraventricular hypothalamus (PVH) from AgRP-IRES-Cre::tdTomato mice under standard diet (SD), or after 1-day or 5-day caloric restriction (CR). Scale bar: 50 µm. **(B–C)** Quantification of AgRP fiber density (**B**) and tdTomato fluorescence intensity (**C**) in anterior, mid, and posterior PVH subregions (100-μm intervals). A one-way ANOVA followed by Tukey’s multiple comparisons test revealed a statistically significant increase in mid-PVH fiber density after 1-day CR (*n* = 3 per group). **(D)** Schematic of the optogenetic setup. AAV-FLEX-ChR2-eYFP (or control AAV-FLEX-eYFP) was injected into the arcuate nucleus (ARC) of AgRP-IRES-Cre mice; an optic fiber was implanted 0.2 mm above the ARC. **(E)** Experimental timeline. Mice received photostimulation before and after 1-day or 5-day CR. **(F–G)** One-hour food intake during optogenetic stimulation at 0, 5, 10, and 20 Hz in 1-day (**F**, *n* = 8) and 5-day (**G**, *n* = 5) CR paradigms. Error bars represent SEM. * *p* < 0.05.

AgRP neurons were stimulated at increasing frequencies (0, 5, 10, and 20 Hz), with each frequency applied for 1 hour, followed by a 30-minute off period to minimize behavioral habituation, phototoxicity, and potential channelrhodopsin inactivation from prolonged stimulation. This stepwise stimulation paradigm was designed to capture subtle shifts in feeding behavior, enabling detection of potentiation or depression across the physiological range [8,34]. Despite increased AgRP→PVH innervation, frequency–response curves were unchanged pre- vs post-CR, indicating **state-dependent decoupling** of structure and output (Figure 1F–G).

### High-fat diet reduces AgRP→PVH innervation but enhances orexigenic output

Having shown that caloric restriction increases AgRP→PVH innervation without amplifying the feeding response, we next asked whether the reciprocal metabolic state, short-term caloric excess, would produce the opposite structural and functional effects. Changes in energy availability have been linked to remodeling of AgRP→PVH connectivity, with some evidence that high-fat diet (HFD) may lead to diminished innervation of this downstream target [18].

To mirror our earlier CR experiments, we assessed AgRP fiber density in AgRP-IRES-Cre::tdTomato mice following 1-day or 5-day HFD, quantifying projections across anterior, middle, and posterior PVH subregions. Similar to the CR condition, changes were localized to the mid-PVH. We observed a significant reduction in AgRP fiber density and tdTomato signal intensity after 5-day HFD, with no difference in anterior or posterior PVH (Figure 2A–C). No reduction was seen after 1-day HFD, suggesting that structural retraction requires sustained exposure to high calorie food.

**Figure 2.**
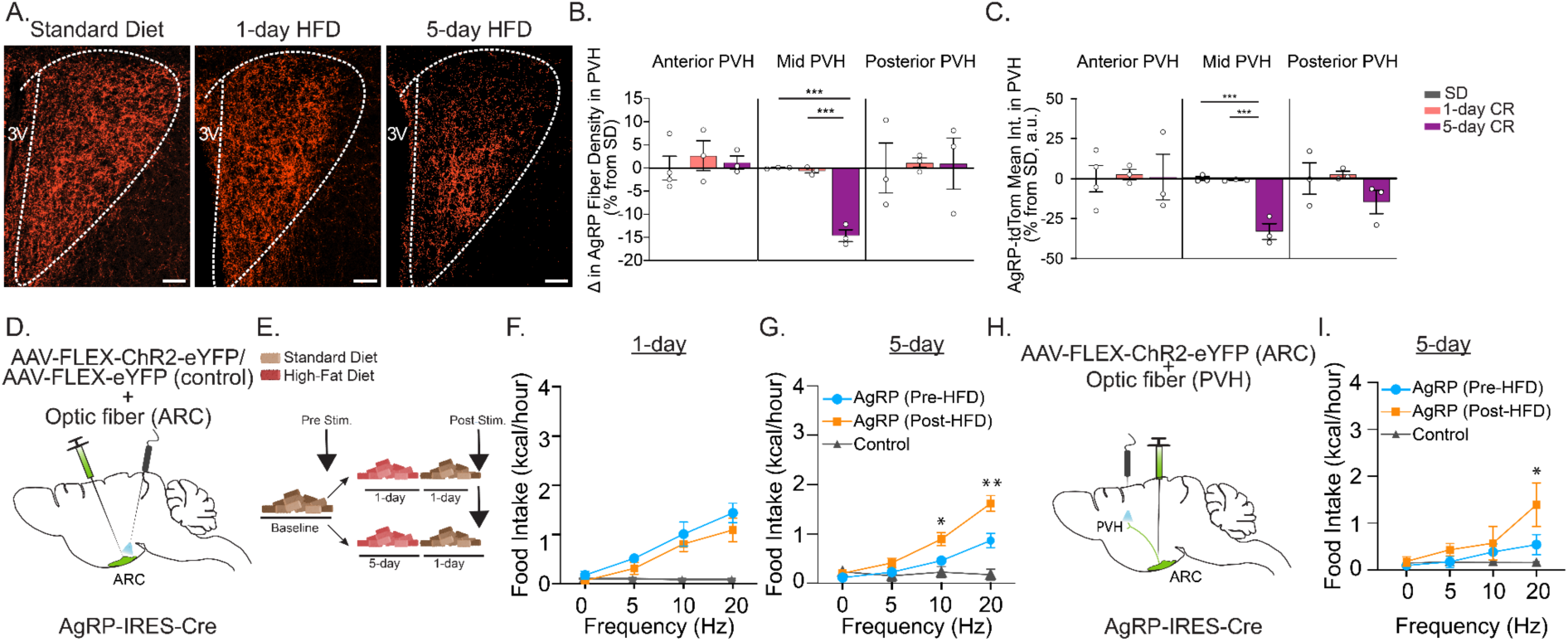
High-fat diet remodels AgRP→PVH innervation and enhances AgRP neuron-driven food intake. (A) Representative images showing AgRP fiber labeling (tdTomato) in the PVH under SD, 1-day, or 5-day HFD. Scale bars: 50 μm. (B–C) Quantification of AgRP fiber density (B) and tdTomato fluorescence intensity (C) in anterior, mid, and posterior PVH subregions, expressed as percent change relative to SD. N = 3 per group. One-way ANOVA with Tukey’s multiple comparisons test. (D) Schematic of viral injection and optic fiber placement for AgRP soma stimulation in the ARC. (E) Experimental timeline for optogenetic testing before and after dietary manipulation. (F–G) Frequency-dependent food intake in response to AgRP soma stimulation following 1-day (F, N = 5) or 5-day HFD (G, N = 11); control group N = 3. (H) Schematic of PVH terminal stimulation. (I) Food intake in response to PVH terminal stimulation after 5-day HFD (N = 6). Two-way ANOVA with Tukey’s multiple comparisons test was used for all optogenetic experiments. Error bars represent SEM. *p < 0.05, **p < 0.01, ***p < 0.001.

**Supplementary Figure 1.**
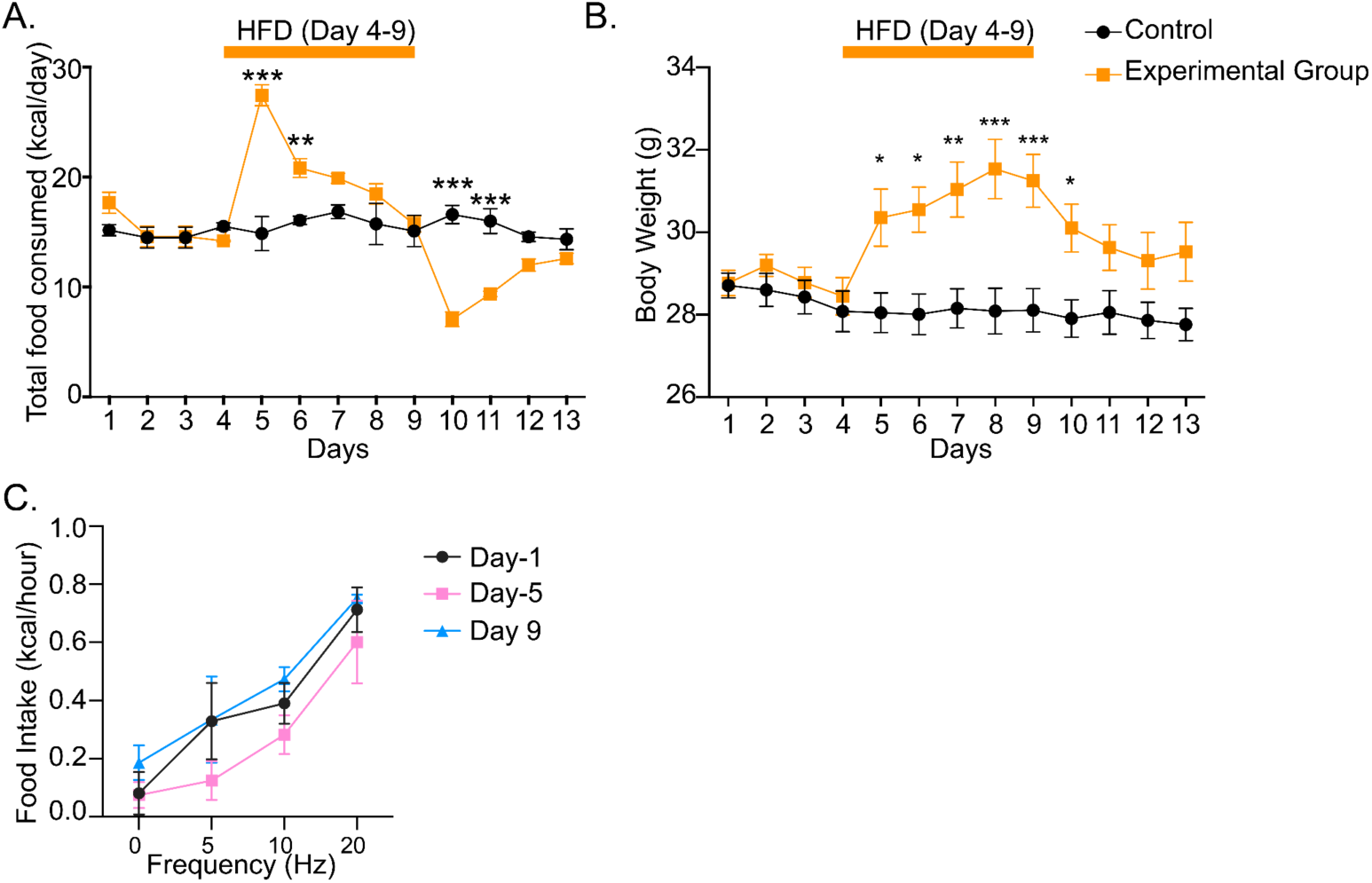
Daily caloric intake, body weight, and AgRP-driven feeding during a short-term HFD paradigm. (A) Daily total food consumption (kcal/day) in control and experimental groups across 13 days. Animals were maintained on standard chow except during days 4–9, when they were provided high-fat diet (HFD). (B) Daily body weight (g) of the same groups during the 13-day period, with HFD exposure from days 4–9. (C) Food intake (kcal/hour) measured during optogenetic stimulation at 1, 5, 10, and 20 Hz on day 1, day 5, and day 9. Data are presented as mean ± SEM. *p < 0.05, **p < 0.01, ***p < 0.001.

We then asked whether this anatomical remodeling altered AgRP functional output. As in the CR paradigm, AgRP-IRES-Cre mice expressing ChR2 received optogenetic stimulation before and after HFD exposure (Figure 2D–E). To control for SD devaluation effects known to follow short-term HFD [35,36], animals were returned to standard chow for 24 hours prior to testing.

Under all conditions, AgRP stimulation induced a frequency-dependent increase in food intake. However, following 5-day HFD, AgRP activation elicited significantly greater feeding compared to baseline, despite the reduction in PVH innervation (Figure 2F–G). This increased orexigenic output was also observed when AgRP terminals in the PVH were directly stimulated (Figure 2H–I).

Daily caloric intake and body weight were transiently elevated during HFD access and returned to baseline after the switch to standard diet, confirming the effectiveness and reversibility of the dietary manipulation (Supplementary Figure 1A–B). A significant reduction in feeding was observed across the first two days following the switch from HFD to standard chow, consistent with SD devaluation. On the first day of HFD exposure, AgRP-driven feeding was modestly reduced, though not significantly, possibly reflecting a transient sensory or motivational shift (Supplementary Figure 1C).

Taken together, these findings reveal that short-term HFD leads to structural retraction of AgRP→PVH projections while enhancing AgRP-driven feeding. In parallel with our CR findings, these results underscore that the animal’s physiological state, rather than projection density alone, determines the functional efficacy of AgRP neurons within hypothalamic feeding circuits.

### BDNF expression in the PVH increases in response to both caloric restriction and high-fat diet

After discovering that 5-day HFD amplifies the orexigenic impact of AgRP neurons despite reduced AgRP→PVH innervation, we reasoned that a postsynaptic gain mechanism within the PVH might account for this heightened responsiveness. BDNF emerged as a strong candidate: (i) it is a well-established mediator of activity-dependent synaptic plasticity and neuronal remodeling across many brain regions [20,37–42], and (ii) its expression in the PVH has been directly linked to the regulation of feeding behavior and energy homeostasis [25,43,44]. Consistent with this, re-analysis of single-cell transcriptomic data [53] revealed broad BDNF expression across PVH neuronal clusters (Figure 3A).

**Figure 3.**
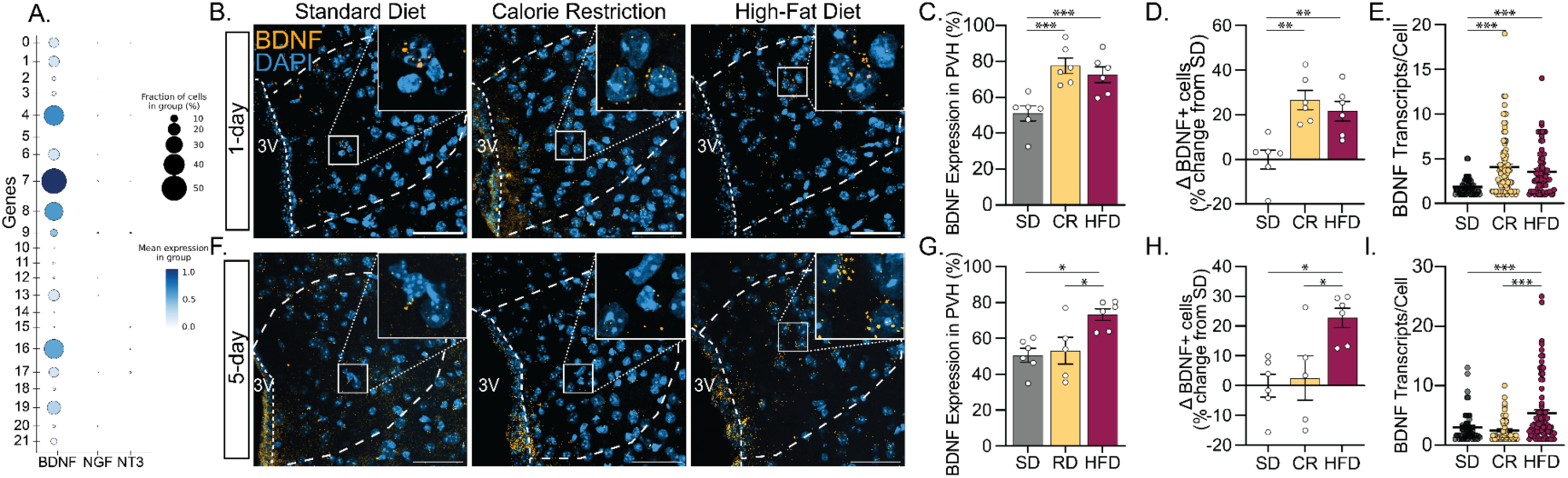
BDNF expression in the PVH is dynamically regulated by acute and sustained metabolic challenges. (A) Dot plot showing expression of neurotrophins (BDNF, NGF, NT3) across PVH neuronal clusters, based on single-cell RNA-sequencing data from the mouse HypoMap [53]. (B, F) Representative RNAscope images showing BDNF mRNA (yellow puncta) in PVH sections from mice maintained on standard diet (SD), or after 1-day caloric restriction (CR), 1-day high-fat diet (HFD) (B), and 5-day CR or 5-day HFD (F). Scale bars: 50 μm. (C, G) Quantification of the percentage of BDNF-expressing cells in the PVH across dietary conditions. (D, H) Percent change in BDNF-expressing cells relative to SD controls. (E, I) Quantification of BDNF transcript abundance per cell. N = 3 per group. One-way ANOVA followed by Tukey’s multiple comparisons test. Error bars represent SEM. *p < 0.05, **p < 0.01, ***p < 0.001.

We therefore asked whether BDNF expression in the PVH is dynamically regulated by changes in metabolic state. Using RNAscope in situ hybridization, we quantified BDNF mRNA in PVH sections from mice subjected to 1-day CR or 5-day HFD, alongside SD controls. Surprisingly, both 1-day CR and 1-day HFD significantly increased the percentage of BDNF-expressing cells and the number of BDNF transcripts per cell relative to SD (Figure 3B–E). We next examined whether these effects were sustained with prolonged dietary interventions. While 5-day CR no longer elevated BDNF expression, 5-day HFD continued to significantly increase both the percentage of BDNF-positive cells and the transcript abundance per cell (Figure 3F–I). These results indicate that BDNF expression in the PVH is rapidly upregulated by acute shifts in energy state but remains persistently elevated only under sustained caloric excess.

### Reciprocal regulation of BDNF receptors in AgRP neurons under opposing energetic states

We found that BDNF expression in the PVH increases under both CR and HFD conditions, indicating that BDNF availability alone cannot account for the different AgRP circuit changes observed. BDNF is known to signal through two receptors with distinct roles: TrkB, which is typically associated with neuronal growth and synaptic stabilization [45,46], and p75NTR, which is often linked to synaptic pruning and structural remodeling [47]. This duality raises the possibility that BDNF functions as a permissive signal for neuronal plasticity, with the direction of adaptation determined by differential receptor engagement rather than by BDNF itself acting as an instructive cue. We therefore hypothesized that AgRP neurons undergo state-dependent shifts in receptor expression that shape their response to BDNF under different metabolic conditions.

To test this, we first confirmed that AgRP neurons express both BDNF receptors under basal conditions. Using RNAscope in situ hybridization, we detected TrkB and p75NTR transcripts in AgRP neurons of mice maintained on SD, demonstrating that these neurons are equipped to respond to BDNF through both signaling pathways (Figure 4A,D).

**Figure 4.**
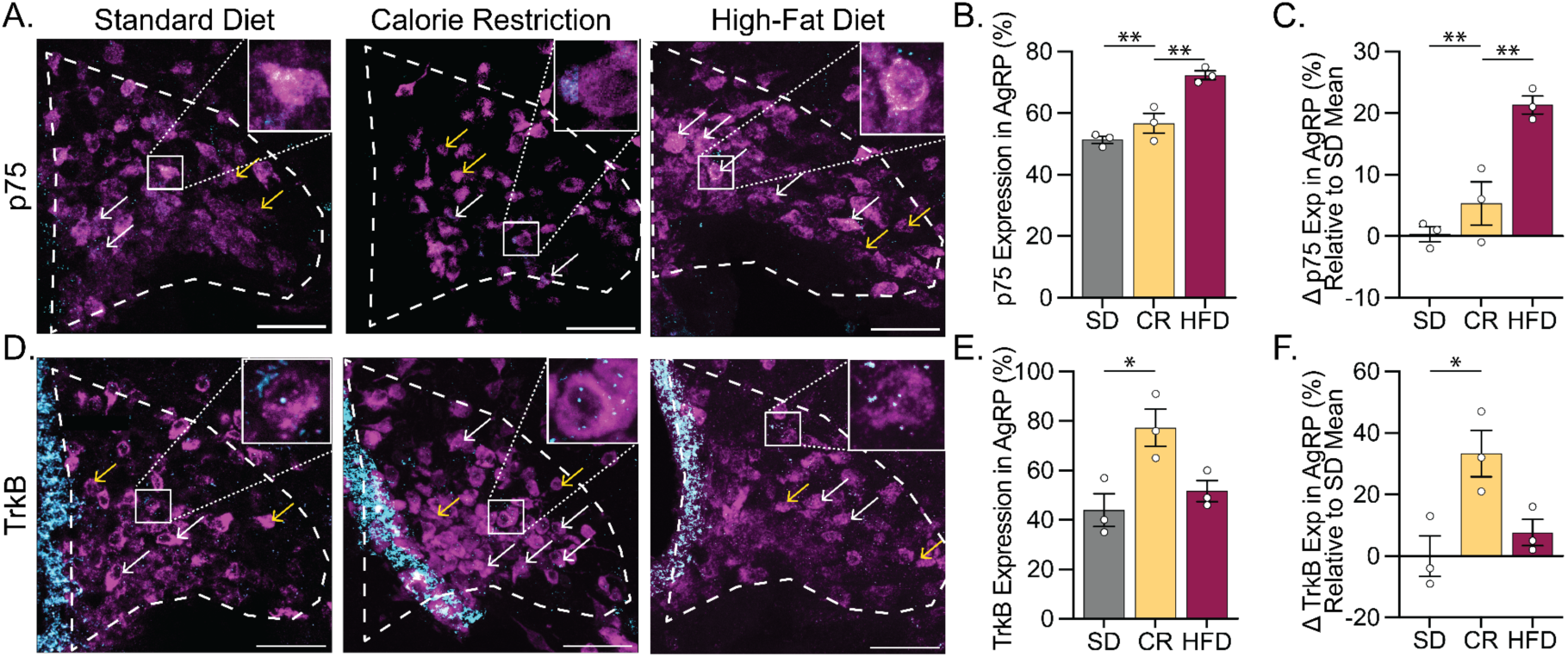
Caloric state bidirectionally regulates BDNF receptor expression in AgRP neurons. (A) Representative RNAscope fluorescent in situ hybridization images showing p75NTR mRNA expression (cyan) in AgRP neurons (magenta) within the ARC under SD, CR, or HFD conditions. Insets highlight examples of AgRP neurons expressing p75NTR (white arrows) and non-expressing AgRP neurons (yellow arrows). Scale bars, 50 µm. (B) Quantification of the percentage of AgRP neurons expressing p75NTR across dietary groups. (C) Relative change in p75NTR expression, normalized to SD group mean. (D) Representative RNAscope images showing TrkB mRNA (cyan) in AgRP neurons (magenta) under SD, CR, or HFD conditions, with inset annotations as in (A). (E) Quantification of the percentage of AgRP neurons expressing TrkB across conditions. (F) Relative change in TrkB expression, normalized to SD group mean. Data are presented as mean ± SEM; each dot represents an individual animal. *p < 0.05, **p < 0.01; one-way ANOVA with Tukey’s multiple comparisons test.

We next assessed whether caloric challenges alter receptor expression in AgRP neurons. RNAscope analysis revealed a distinct pattern of reciprocal regulation. Following 5 days of HFD, the percentage of AgRP neurons expressing p75NTR increased significantly relative to SD controls (Figure 4A–C), whereas TrkB expression remained unchanged. In contrast, 24-hour CR selectively upregulated TrkB expression in AgRP neurons (Figure 4D–F) without affecting p75NTR levels.

These findings confirm that AgRP neurons exhibit dynamic, bidirectional regulation of BDNF receptor expression in response to caloric challenges, with p75NTR upregulated during HFD and TrkB during CR. This receptor-specific shift drives opposing structural and functional adaptations in AgRP circuits, revealing a molecular rheostat that fine-tunes hunger signaling to metabolic context and highlighting potential targets for modulating energy homeostasis.

### BDNF Receptors Differentially Regulate Baseline AgRP→PVH Innervation

Given that caloric state alters both AgRP→PVH innervation and BDNF receptor expression in AgRP neurons, we next tested whether these receptors directly regulate AgRP neuron projections. Specifically, we asked whether TrkB or p75NTR are required to maintain baseline AgRP→PVH innervation under standard diet conditions. To address this, we analyzed AgRP fiber density and projection intensity in the PVH of AgRP-TrkB-KO and AgRPp75-KO mice maintained on SD.

AgRPp75-KO mice displayed a significant increase in AgRP fiber density and projection intensity relative to controls under SD conditions (p < 0.01; figure 5A–C). This finding suggests that p75NTR constrains AgRP neuron innervation under basal conditions, likely by limiting axonal growth or promoting synaptic pruning.

**Figure 5.**
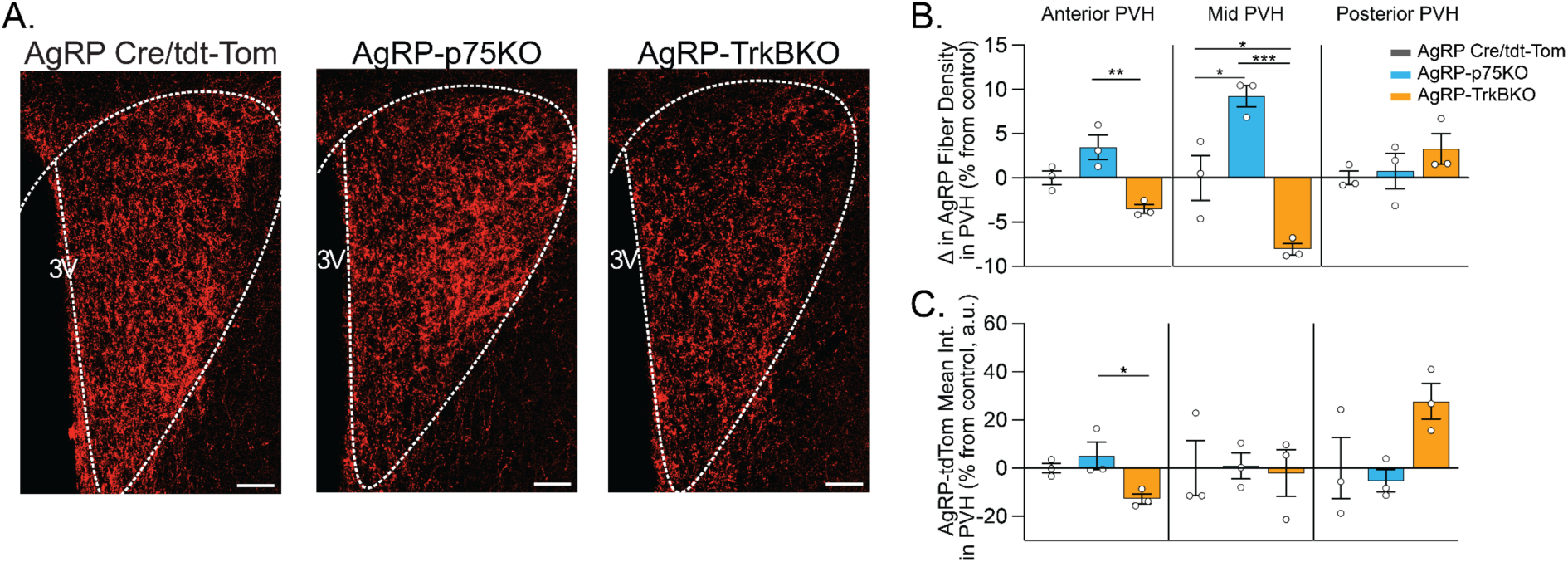
BDNF receptors differentially regulate AgRP→PVH innervation. (A) Representative images showing AgRP-tdTomato-labeled fibers in the PVH of control (AgRP-IRES-Cre::tdTomato), AgRP-specific TrkB knockout (AgRP-TrkBKO), and p75 knockout (AgRP-p75KO) mice. Scale bars: 50 μm. (B) Quantification of AgRP fiber density across anterior, mid, and posterior PVH subregions, expressed as percent change from control. (C) Mean AgRP-tdTomato fluorescence intensity across PVH subregions. N = 3 per group. One-way ANOVA followed by Tukey’s multiple comparisons test. Error bars represent SEM. *p < 0.05, **p < 0.01, ***p < 0.001.

In contrast, AgRP-TrkB-KO mice exhibited a significant reduction in AgRP fiber density and projection intensity within the PVH compared to littermate controls (p < 0.01; figure 5D–F). This result indicates that TrkB is required for the maintenance of normal AgRP→PVH innervation under baseline conditions, consistent with its role in supporting neuronal growth and synaptic stabilization.

Together, these results demonstrate that TrkB and p75NTR exert opposing influences on AgRP→PVH connectivity even in the absence of external metabolic challenges. TrkB promotes the maintenance of AgRP projections, whereas p75NTR acts to restrict their growth. These findings provide direct evidence that BDNF receptor signaling governs the structural organization of AgRP neurons under basal conditions. Notably, while BDNF itself appears to act as a permissive signal for plasticity, the receptors function as instructive elements that direct the specific structural outcomes.

### BDNF Receptors Differentially Shape the Orexigenic Response of AgRP Neurons under Caloric Challenges

Having established that TrkB and p75NTR exert opposing effects on AgRP neuron innervation, we next tested whether these receptors modulate the functional output of AgRP neurons under metabolic challenges. To document the dietary manipulations in these cohorts, daily food intake and body weight were tracked during the 5-day HFD protocol (Supplementary Figure 2). We assessed optogenetically-evoked food intake in AgRP-TrkBKO and AgRP-p75KO mice before and after 1-day CR or 5-day HFD.

Under CR conditions, AgRP-p75KO mice displayed minimal feeding responses to AgRP neuron stimulation, with no enhancement following CR (Figure 6A). In contrast, AgRP-TrkBKO mice exhibited robust, frequency-dependent increases in food intake both before and after CR, though CR itself did not further potentiate feeding responses (Figure 6B). A similar pattern emerged under HFD conditions: AgRP-p75KO mice showed persistently blunted feeding, while AgRP-TrkBKO mice maintained strong orexigenic responses to AgRP stimulation, with no difference between pre- and post-HFD conditions (Figure 6E, F).

**Figure 6.**
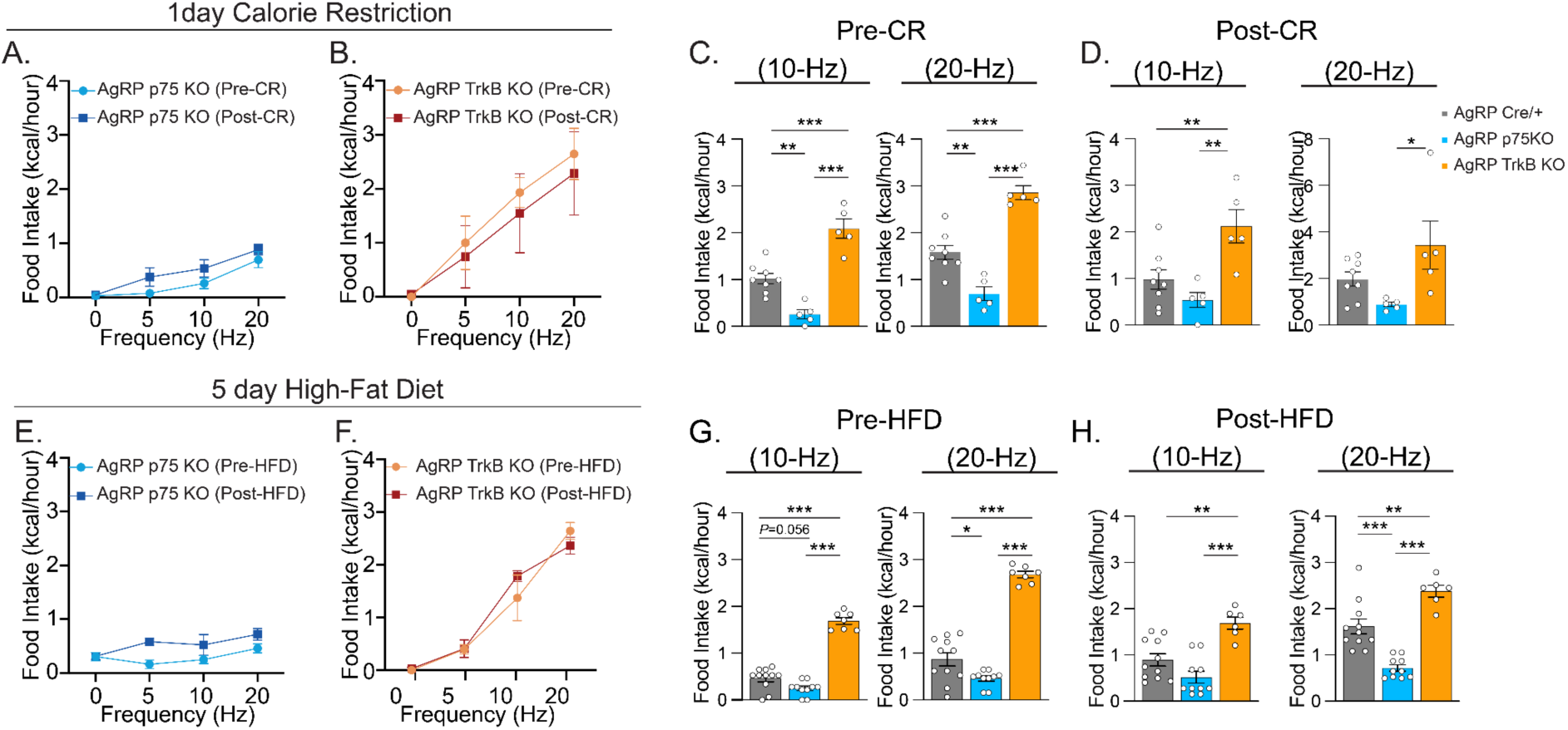
TrkB and p75NTR differentially regulate feeding responses to AgRP neuron activation under caloric restriction and high-fat diet. (A, B) Optogenetically-evoked food intake in AgRP-p75KO (A) and AgRP-TrkBKO (B) mice before and after 1-day CR. (C, D) Quantification of food intake at 10 Hz and 20 Hz stimulation frequencies before (C) and after (D) CR. (E, F) Optogenetically-evoked food intake in AgRP-p75KO (E) and AgRP-TrkBKO (F) mice before and after 5-day high-fat diet (HFD). (G, H) Quantification of food intake at 10 Hz and 20 Hz stimulation frequencies before (G) and after (H) HFD. Control data were re-plotted from Figures 1 and 2 to allow direct comparison. Data represent mean ± SEM; each point represents an individual animal. Statistical comparisons were performed using one-way ANOVA followed by Tukey’s post hoc test for genotype comparisons at each frequency and time point. *p < 0.05, **p < 0.01, ***p < 0.001.

**Supplementary Figure 2.**
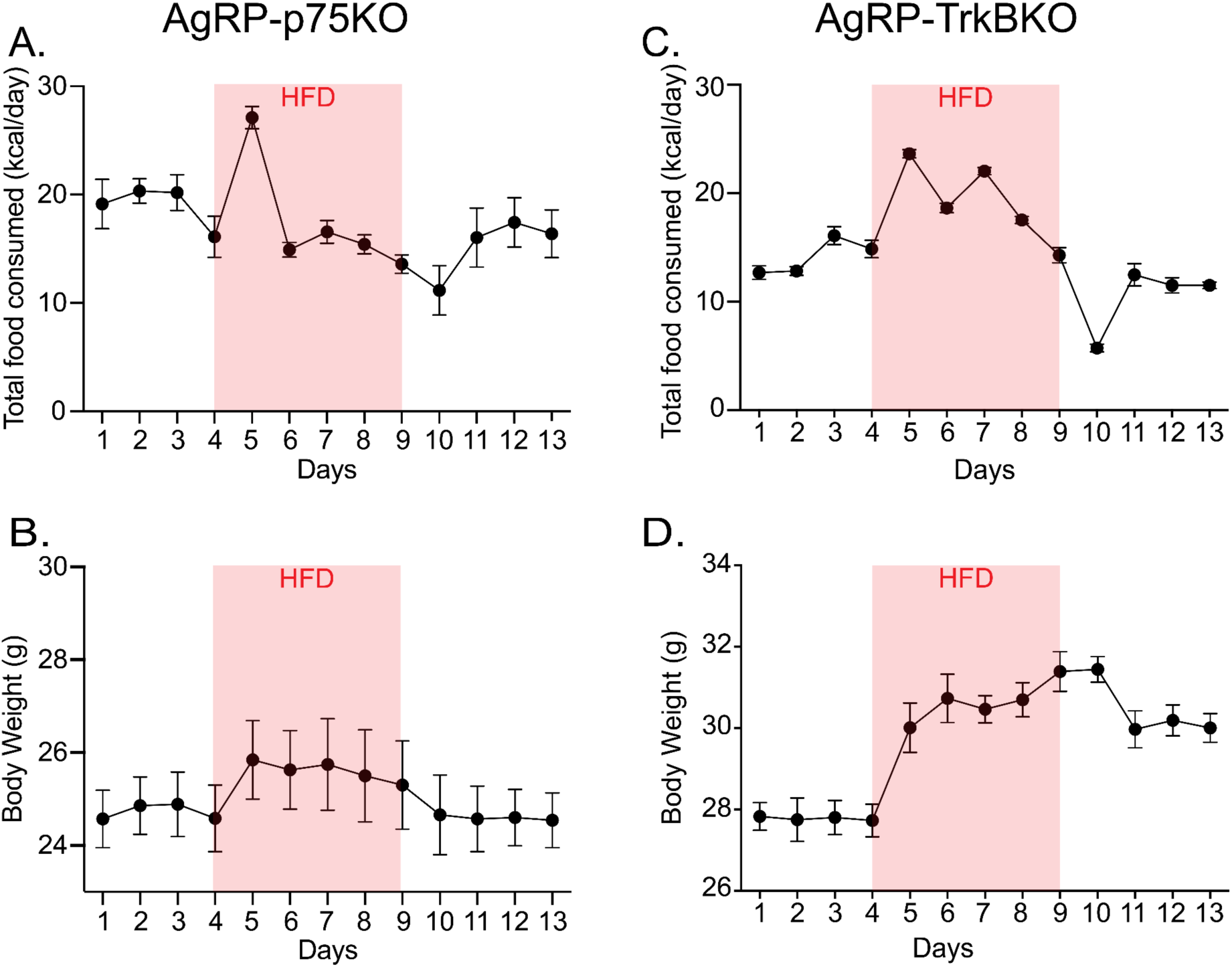
Daily food intake and body weight trajectories in AgRP neuron-specific p75NTR and TrkB knockout mice during high-fat diet exposure. (A–B) Daily food intake (A) and body weight (B) measurements in AgRP-p75KO mice across a 13-day paradigm including a 5-day high-fat diet (HFD; shaded pink). (C–D) Daily food intake (C) and body weight (D) measurements in AgRP-TrkBKO mice across the same HFD paradigm. Data are shown as mean ± SEM.

However, a striking divergence became evident when the KO models were directly compared to AgRP-Cre/+ controls. Across both CR and HFD paradigms, AgRP-p75KO mice consistently exhibited significantly lower feeding responses at both 10 Hz and 20 Hz stimulation frequencies, while AgRP-TrkBKO mice showed significantly elevated food intake compared to controls (Figure 6C, D for CR; Figure 6G, H for HFD).

These results demonstrate that TrkB and p75NTR regulate the orexigenic capacity of AgRP neurons in distinct and opposing ways. Neither CR nor HFD altered stimulation-driven feeding within a given genotype, yet the knockout phenotypes reveal opposing baseline states: AgRP-p75KO mice exhibited persistently reduced responses, likely reflecting a floor effect from elevated basal innervation, while AgRP-TrkBKO mice showed exaggerated feeding responses, consistent with reduced basal innervation that preserves greater stimulation capacity. These opposing phenotypes suggest that the receptor knockouts effectively lock the system into extreme configurations, eliminating the dynamic range necessary for adaptive plasticity. In contrast to wild-type circuits that can dissociate structure and function depending on metabolic state, the knockouts remain fixed in their respective positions, unable to further adjust in response to caloric challenges. This pattern indicates that balanced BDNF receptor signaling is essential for the flexible control of AgRP→PVH connectivity and the circuit’s potential for state-dependent adaptation.

## Discussion

Hypothalamic feeding circuits must adapt to fluctuating energy conditions, yet the molecular mechanisms enabling this flexibility remain poorly understood. Our findings reveal an unexpected principle: AgRP neuron projections to the PVH and their functional output can be dissociated depending on metabolic state, challenging conventional assumptions about how connectivity determines behavior in homeostatic circuits. This dissociation emerges through a molecular rheostat mechanism involving bidirectional regulation of BDNF receptors in AgRP neurons, where the balance between TrkB and p75NTR expression dictates both the structural organization and functional capacity of the circuit.

We demonstrate that caloric restriction increases AgRP→PVH fiber density within 24 hours, particularly in mid-PVH regions, consistent with prior reports of fasting-induced structural remodeling[16,17]. Yet despite this anatomical enhancement, optogenetic stimulation of AgRP neurons evoked similar feeding responses before and after restriction. This dissociation between increased fiber density and unchanged functional output occurs alongside upregulated TrkB expression in AgRP neurons. TrkB signaling is known to promote axonal growth through PI3K-Akt pathways[31] and can modulate neuronal excitability through effects on potassium channels [48,49], though the specific mechanisms mediating this structure-function dissociation remain to be determined.

The response to caloric excess presents a striking counterpoint. Five days of high-fat diet reduced AgRP→PVH innervation, aligning with observations by Wei et al.[18], yet paradoxically amplified feeding responses to both somatic and terminal stimulation. This apparent contradiction resolves when viewed through the lens of receptor-mediated plasticity: HFD selectively upregulates p75NTR in AgRP neurons while sustaining elevated BDNF expression in the PVH. The p75NTR receptor, traditionally associated with axonal pruning [50], can simultaneously enhance neuronal excitability[51], providing a mechanistic basis for how circuits can reduce connectivity while potentiating output. This dual action effectively ‘turns the rheostat’ to maximize feeding drive during caloric abundance, potentially an evolutionary strategy to promote energy storage when resources are plentiful.

The receptor knockout phenotypes provide crucial validation of this model while revealing its dynamic range. AgRP neurons lacking p75NTR displayed increased basal innervation but blunted feeding responses, whereas TrkB deletion produced the opposite pattern: reduced innervation with exaggerated feeding. Critically, neither knockout could adapt to metabolic challenges, CR and HFD failed to modulate their responses, suggesting these manipulations effectively ‘break the dial’ by fixing it at extreme positions. This loss of flexibility underscores that balanced BDNF signaling is essential not just for baseline function but for the adaptive capacity that defines healthy metabolic regulation.

The regional specificity of these changes to mid-PVH warrants consideration. This subregion may express unique combinations of BDNF receptors or downstream effectors that make it particularly sensitive to neurotrophin-mediated remodeling. Alternatively, mid-PVH neurons might represent a functionally specialized population that serves as the primary integration site for AgRP-derived hunger signals. The rapid timeline of CR-induced changes (peaking at 1 day) versus the sustained requirement for HFD effects (5 days) further suggests distinct activation thresholds or kinetics for TrkB versus p75NTR signaling pathways, though these temporal dynamics require further investigation.

Our findings extend beyond simple activity-dependent plasticity to reveal an adaptation where circuits adjust their properties to match metabolic context rather than merely responding to use. The incomplete functional modulation under CR, where structure changes without altering output, points to additional regulatory inputs that remain to be identified. These might include ghrelin-independent hormonal pathways, duration-dependent signals beyond our 1-5 day timeframe, or competing synaptic inputs that constrain AgRP neuron efficacy.

This work also connects to broader principles of circuit organization. The dissociation between structure and function we document in feeding circuits may extend to other brain regions where BDNF regulates both anatomy and physiology. Recent studies have highlighted similar paradoxes in other hypothalamic populations[25,26], suggesting that neurotrophin-mediated uncoupling of connectivity and output might be a general feature of homeostatic systems. The concept of receptor balance as a molecular rheostat could thus provide a unifying framework for understanding how neural circuits maintain flexibility while preserving core functions.

Several limitations constrain our interpretation. The short time windows we examined for CR and HFD may not capture the chronic adaptations that accompany sustained metabolic challenges or obesity. Our use of constitutive AgRP-specific deletions could introduce developmental compensation that obscures the acute roles of these receptors; future studies employing inducible, adult-restricted manipulations will be essential to address this concern. While RNAscope allowed us to quantify Bdnf transcript levels, this approach cannot distinguish between proBDNF and mature BDNF forms, a critical distinction given that these ligands can differentially engage p75NTR versus TrkB signaling pathways. Finally, our analysis focused exclusively on the PVH as a downstream target, yet AgRP neurons project to multiple brain regions including the lateral hypothalamus, bed nucleus of the stria terminalis, and paraventricular thalamus, each of which may exhibit distinct state-dependent plasticity rules.

Future work should address whether structural changes precede or follow functional adaptations, potentially using in vivo imaging to capture the temporal evolution of plasticity. Investigating how other AgRP projection targets respond to similar manipulations could reveal whether the PVH represents a unique site of regulation or exemplifies a broader principle. The incomplete engagement of the rheostat under CR particularly demands attention identifying the missing signals that would fully activate orexigenic drive during energy deficit could illuminate new regulatory pathways. Inducible genetic approaches combined with longer dietary interventions will help establish whether the plasticity mechanisms we identify contribute to the persistent hunger following weight loss[52] or the rapid weight regain that challenges clinical interventions.

By demonstrating that metabolic state determines the relationship between circuit structure and function through receptor-specific neurotrophin signaling, our findings challenge the traditional view that anatomical connectivity directly predicts behavioral output. Instead, we reveal that hypothalamic feeding circuits employ a sophisticated molecular rheostat that enables context-appropriate adaptations, preparing for opportunity during scarcity while maximizing intake during abundance. This framework not only advances our understanding of homeostatic plasticity but also suggests that targeting the balance between TrkB and p75NTR signaling, rather than individual pathways, may offer more nuanced approaches for modulating appetite and energy balance.

## Materials and Methods

### Mouse Lines, Breeding, and Husbandry

All experiments were conducted in accordance with protocols approved by the University of Virginia Institutional Animal Care and Use Committee and in compliance with the Association for Assessment and Accreditation of Laboratory Animal Care (AAALAC) policies. Mice were housed in temperature-controlled rooms (22 ± 1°C) under a 12:12-hour light/dark (LD) cycle (lights on at 6:00 AM), with ad libitum access to food (PicoLab Rodent Diet 5053) and water unless otherwise specified. Male and female mice aged 8 weeks or older were used; no a priori sex effects were observed, so data were pooled across sex. Data from both sexes were analyzed independently and combined when no significant sex differences were observed. The following transgenic mouse lines were used: AgRP-ires-Cre (Agrptm1(cre)Lowl/J, JAX #012899), p75NTR-floxed (Ngfrtm1.1Vk/Bnap/J, JAX #031162), TrkB-floxed (Ntrk2tm1Ddg/J, JAX #022363), and Rosa26-LSL-tdTomato (Ai9; B6.Cg-Gt(ROSA)26Sortm9(CAG-tdTomato)Hze/J, JAX #007909). AgRP-specific knockout lines referred to as AgRP-p75KO and AgRP-TrkBKO were generated by crossing AgRP-Cre mice with p75NTR-floxed or TrkB-floxed animals, respectively. For anatomical tracing and fiber density analyses, AgRP-Cre mice were crossed with Ai9 reporter mice to label AgRP neurons with tdTomato fluorescence. All mouse lines were maintained on a C57BL/6J background and backcrossed for at least five generations prior to experimentation.

### Dietary Interventions

Mice were assigned to one of three dietary conditions. In the standard diet (SD) condition, animals were maintained on PicoLab® Rodent Diet 20 (5053), which provides approximately 21% protein and 5% fat by weight, with an energy density of ∼3.4 kcal/g. Food and water were available ad libitum throughout the experiment. For caloric restriction (CR), food was removed for 24 hours starting at zeitgeber time 0 (6:00 AM, lights on), while water remained available ad libitum. After the restriction period, animals were returned to standard diet for 24 hours before optogenetic testing to minimize the acute effects of deprivation; however, for innervation experiments, animals were sacrificed immediately after the CR period. For the high-fat diet (HFD) condition, mice received ad libitum access to Open Source Diet D12451 (Research Diets), which provides 45% kcal from fat and 20% from protein, with an energy density of 4.73 kcal/g, for five consecutive days. Similar to the CR group, HFD animals were returned to standard diet for 24 hours before optogenetic testing. This 24-hour standard diet recovery period was used to normalize feeding behavior and minimize confounding effects of acute dietary manipulations across groups.

### Stereotaxic Surgeries

#### Viral Injections and Optical Fiber Implantation

Mice were anesthetized with isoflurane (5% induction and 2–2.5% maintenance; Isothesia) and placed in a stereotaxic apparatus (AWD). A heating pad was used throughout surgery to maintain body temperature, and ocular lubricant was applied to prevent corneal desiccation.

For optogenetic experiments, a total of 200–400 nL per side of either pAAV1-EF1a-DIO-hChR2(H134R)-EYFP-WPRE-HGHpA (plasmid from Addgene, #20298; RRID:Addgene_20298; UNC Vector Core, titer ≥ 7 × 10¹² vg/mL) or control virus pAAV1-EF1a-DIO-EYFP (plasmid from Addgene, #27056; RRID:Addgene_27056; UNC Vector Core, titer ≥ 1 × 10¹³ vg/mL) was microinjected unilaterally into the arcuate nucleus (ARC). Stereotaxic coordinates relative to Bregma were: mediolateral (ML): ±0.20 mm, anteroposterior (AP): −1.40 mm, dorsoventral (DV): −5.90 mm. Viral infusions were delivered using a glass pipette attached to a microsyringe pump controller (World Precision Instruments, Micro 4) at a rate of 100 nL/min. The pipette was left in place for 10 min post-infusion and fully withdrawn 15–20 min after viral delivery. Following viral injections, fiber optic cannulae (RWD Life Science; Ø1.25 mm ceramic ferrule, 400 μm core, 0.50 NA, length 6 mm; (Cat No: R-FOC-BF400C-50NA) were implanted and secured with dental cement (C&B Metabond, Parkell). For ARC-targeted experiments, the cannula was placed 0.2 mm dorsal to the injection site. For PVH-targeted experiments, viral injections were made in the ARC as above, and the optic fiber was implanted above the PVH at the following stereotaxic coordinates: ML: ±0.20 mm, AP: −0.82 mm, DV: −4.70 mm. Mice were allowed a minimum of 3 weeks recovery to ensure adequate opsin expression before behavioral testing. All surgical procedures were performed under sterile conditions and in accordance with institutional animal care and use guidelines. Animals with mistargeted injections or misplaced fiber implants were excluded from subsequent analyses.

### Optogenetic Stimulation and Feeding Behavior

Mice were habituated for 5 consecutive days to handling, the experimental chamber, the patch cord, and the presence of standard chow (SD). Habituation included mock stimulation sessions (0 Hz, light off) to control for cord attachment and tethering effects. On test days, food was always SD, and animals were not food-deprived prior to experiments (ad libitum access maintained). For stimulation, a 473 nm laser delivered 10 ms pulses at 0–20 Hz (8–10 mW at the fiber tip, calibrated with a power meter; Thorlabs). Light was delivered in 1 s on / 3 s off trains. Frequencies were tested in ascending order within the same session, with 30 min interleaved intervals between stimulation blocks. Each stimulation frequency was applied for 1 h, during which food was available ad libitum. Food intake was measured by weighing pre-measured chow pellets immediately before and after each stimulation period. Intake was expressed as grams consumed during the 1-h stimulation block. Food consumption was also measured during the 30-min interleaved intervals (light off) to assess whether effects of optogenetic stimulation persisted beyond the stimulation period. Experiments were conducted between ZT3–ZT9 (lights on at ZT0 = 6:00 AM). Only mice with confirmed unilateral ARC targeting and correct fiber placement were included; those with mistargeted injections or misplaced fibers were excluded from analysis.

### RNAscope Fluorescence In Situ Hybridization

Brains were dissected from C57BL/6J mice, fixed in 4% paraformaldehyde (PFA) in PBS for 24 h at 4°C, and subsequently cryoprotected in 30% sucrose for 72 h. Fixed tissue was embedded in OCT, frozen on dry ice, and stored at −80°C until sectioning. Coronal sections were cut at 15 µm thickness using a cryostat and mounted onto positively charged slides (Superfrost Plus, Thermo Fisher Scientific, 6776214), then air dried overnight in the dark. The following day, slides were washed twice in PBS for 2 min each, incubated with RNAscope hydrogen peroxide solution (Advanced Cell Diagnostics, 322381) for 10 min at room temperature, rinsed twice in distilled water, and treated with Protease IV (ACD, 322381) for 30 min at 40°C in a HybEZ II oven (ACD, 321710/321720). Sections were then hybridized for 2 h at 40°C with RNAscope probes directed against Mm-Agrp (Ref. 400711, Lot 24088A), Mm-Ngfr (Cat. No. 494261), Mm-Ntrk2 (Cat. No. 423611), or Mm-Bdnf-O1 (Cat. No. 461561). Probes were assigned to separate channels to enable multiplex detection. After hybridization, slides were washed twice for 2 min each in 1× RNAscope wash buffer (ACD, 310091) and subjected to signal amplification using the RNAscope Multiplex Fluorescent Detection Kit v2 (ACD, 323110). Amplification steps consisted of sequential incubations in AMP1, AMP2, and AMP3 solutions, each for 30 min at 40°C, with two 2-min washes in 1x wash buffer between steps. Fluorescent detection was performed channel by channel using horseradish peroxidase (HRP) development and tyramide signal amplification. For Channel 1 (Agrp), slides were incubated with HRP-C1 for 15 min at 40°C, washed twice, and developed with TSA Fluorescein (Akoya, TS-000200, Lot 20220305) diluted 1:750 in RNAscope TSA buffer (ACD, 322809) for 30 min at 40°C, followed by HRP blocker for 15 min. The same steps were repeated for Channel 2 (Ngfr) using HRP-C2 and TSA Cy3 (Akoya, TS-000202, Lot 20203437, 1:750), and for Channel 3 (Ntrk2 or Bdnf-O1) using HRP-C3 and TSA Cy5 (Akoya, TS-000203, Lot 20233412, 1:750). Each HRP incubation was followed by two 2-min washes, and each dye incubation by two 2-min washes before proceeding to HRP block. Following signal detection, sections were counterstained with DAPI and mounted with Fluoromount-G (SouthernBiotech, 0100-20). Slides were allowed to cure at room temperature in the dark for at least 30 min prior to imaging.

### Microscopy and Image Analysis

#### Confocal Microscopy

Imaging was performed on a Zeiss LSM 980 NLO equipped with GaAsP detectors. RNAscope sections were acquired with a 40x oil objective and innervation datasets with a 20x objective. Channels were acquired sequentially to minimize spectral bleed-through with the following excitation/detection bands: DAPI, 405 nm / 420–480 nm; Fluorescein, 488 nm / 500–550 nm; Cy3, 561 nm / 570–620 nm; Cy5 or tdTom channel, 633 nm / 640–700 nm. The pinhole was set to 1 Airy Unit for each channel. Images were collected at 1024×1024 pixels unidirectional scan at standard speed with line averaging 2–4x. For z-stacks, optical sections were acquired at 0.5–1.0 µm steps spanning the full thickness of the section. Acquisition parameters (laser power, detector gain/offset, pinhole, scan speed, averaging, pixel size) were optimized on control tissue and then held constant across animals within each experiment. Z-stacks were exported as maximum intensity projections in ZEN; only linear adjustments to brightness and contrast were applied, identically across groups. Bit depth and scaling were preserved for quantification.

### Quantification of AgRP Fiber Density

AgRP fiber innervation in the PVH was quantified in QuPath (v0.4.3). The PVH was identified using anatomical landmarks from the Franklin & Paxinos mouse brain atlas with DAPI counterstaining to delineate nuclear boundaries. To capture region-specific effects while ensuring comparability across animals, the entire PVH was segmented into matched 100-µm rostrocaudal bins (anterior, middle, posterior) defined from identical atlas windows in every animal; the mid-PVH bin corresponded to approximately AP −0.70 to −0.94 mm from bregma. For each section, subregions were manually outlined, confocal z-stacks were converted to maximum-intensity projections, and pixel size was calibrated in QuPath. Within each masked region, fiber density was computed as the percentage of area occupied by tdTomato-positive signal after applying a uniform, cohort-wide threshold (calibrated on control tissue and then fixed across all images); fluorescence intensity was quantified as the mean gray value within the ROI as an index of projection strength. All images were acquired under identical confocal settings (laser power, detector gain, pinhole, scan speed) to enable direct comparisons. For each animal, a minimum of three sections per PVH subregion were analyzed (total 9–12 PVH-containing sections per mouse), and values were averaged across sections to yield a single density and a single intensity measure per subject. Image acquisition and all quantification were performed by an investigator blinded to experimental group. Subregional statistical comparisons are described in Statistical Analysis.

### Paraventricular Hypothalamus scRNA-seq Analysis

Single cell RNA-sequencing data was extracted from the mouse HypoMap [53] in CZ CELLxGENE. Briefly, the HypoMap data set was loaded into Python (v3.11) Jupyter Notebook (v7.4.4.). The paraventricular hypothalamic nucleus was subsetted and reclustered into 22 clusters after normalization, scaling, and principal component analysis (PCA). Genes of interest (ENSMUSG00000048482 [Bdnf], ENSMUSG00000027859 [Ngf], ENSMUSG00000049107 [Ntf3]) were plotted to visualize percent and mean expression level per cluster.

### RNAscope Quantification

RNAscope signal was quantified using QuPath (vQ.4.3.). For BDNF expression in the PVH, nuclei were segmented using DAPI, and the proportion of DAPI-positive cells containing one or more BDNF puncta was calculated. PVH boundaries were defined according to anatomical landmarks from the Franklin & Paxinos mouse brain atlas. For receptor expression analysis in AgRP neurons, AgRP-positive cells were identified based on probe labeling, and transcript puncta corresponding to p75NTR and TrkB were quantified within each cell. Puncta detection was performed using the cell detection module (positive cell detection and subcellular detection) in QuPath, with thresholds for signal intensity and minimum spot size calibrated on control tissue and subsequently applied uniformly across all images. This approach allowed for automated identification of discrete fluorescent puncta while minimizing background noise. At least 30 AgRP neurons per animal were analyzed across multiple sections, and values were averaged to generate one measurement per mouse. All images were acquired under identical confocal settings, and quantification was performed by an investigator blinded to experimental group.

### Statistical Analysis

All data are presented as mean ± SEM unless otherwise specified. Statistical analyses were performed using GraphPad Prism (v10.5.0). Comparisons between two independent groups were analyzed using two-tailed unpaired Student’s t-tests, while within-subject comparisons (e.g., pre- vs. post-dietary intervention) were assessed using paired t-tests. For experiments involving more than two groups, one-way or two-way ANOVA was applied, followed by Tukey’s or Sidak’s post hoc tests as appropriate; repeated measures designs were analyzed using repeated-measures ANOVA with Sidak’s correction. If normality assumptions were not met, non-parametric tests were employed, including Wilcoxon signed-rank tests for paired data, Mann–Whitney U tests for unpaired data, and Kruskal–Wallis tests with Dunn’s post hoc test for multiple group comparisons. Statistical significance was defined as *p* < 0.05, with figures denoting *p* < 0.05 (\**), p < 0.01 (*\*\*), and *p < 0.001 (*\*\*\*). Sample sizes were determined based on preliminary experiments designed to detect at least a 20% effect size with 90% confidence. The specific test used, test statistic, sample size (n), and p-value for each experiment are reported in the figure legends.

## Acknowledgements

We thank the members of the Deppmann, Güler, and Provencio laboratories (University of Virginia) for their helpful comments and suggestions on the preparation of the manuscript. We are also grateful to Stefani A. Mancuso and Shreya R. Nagarajan for technical assistance and help with mouse colony maintenance. We used AI-assisted technologies (OpenAI, Anthropic) as an aid for drafting and grammar proofing the manuscript.

## Author contributions

Conceptualization: OYC, ADG, CDD

Data curation: OYC, SKM, ENG, SSS, MN, SO, KIW, ODL, LV

Formal analysis: OYC, SKM, ENG, MN

Funding acquisition: OYC, ADG, CDD

Investigation: OYC, SKM, ENG, SSS, MN, SO, KIW, ODL, LV, TBG, PRN, MTM, LC

Methodology: OYC, ENG

Project administration: ADG, CDD

Resources: JNC, ADG, CDD

Supervision: OYC, JNC, ADG, CDD

Validation: OYC, ENG, ADG, CDD

Visualization: OYC, with input from ADG, CDD

Writing – original draft: OYC, ADG, CDD

## Bibliography

1. Sternson, S.M. (2013). Hypothalamic survival circuits: blueprints for purposive behaviors. Neuron 77, 810–824.

2. Timper, K., and Brüning, J.C. (2017). Hypothalamic circuits regulating appetite and energy homeostasis: pathways to obesity. Dis. Model. Mech. 10, 679–689.

3. Morton, G.J., Meek, T.H., and Schwartz, M.W. (2014). Neurobiology of food intake in health and disease. Nat. Rev. Neurosci. 15, 367–378.

4. Tran, L.T., Park, S., Kim, S.K., Lee, J.S., Kim, K.W., and Kwon, O. (2022). Hypothalamic control of energy expenditure and thermogenesis. Exp. Mol. Med. 54, 358–369.

5. Gao, Q., and Horvath, T.L. (2008). Neuronal control of energy homeostasis. FEBS Lett. 582, 132–141.

6. Williams, K.W., and Elmquist, J.K. (2012). From neuroanatomy to behavior: central integration of peripheral signals regulating feeding behavior. Nat. Neurosci. 15, 1350–1355.

7. Hahn, T.M., Breininger, J.F., Baskin, D.G., and Schwartz, M.W. (1998). Coexpression of Agrp and NPY in fasting-activated hypothalamic neurons. Nat. Neurosci. 1, 271–272.

8. Aponte, Y., Atasoy, D., and Sternson, S.M. (2011). AGRP neurons are sufficient to orchestrate feeding behavior rapidly and without training. Nat. Neurosci. 14, 351–355.

9. van den Top, M., Lee, K., Whyment, A.D., Blanks, A.M., and Spanswick, D. (2004). Orexigen-sensitive NPY/AgRP pacemaker neurons in the hypothalamic arcuate nucleus. Nat. Neurosci. 7, 493–494.

10. Krashes, M.J., Koda, S., Ye, C., Rogan, S.C., Adams, A.C., Cusher, D.S., Maratos-Flier, E., Roth, B.L., and Lowell, B.B. (2011). Rapid, reversible activation of AgRP neurons drives feeding behavior in mice. J. Clin. Invest. 121, 1424–1428.

11. Sternson, S.M., and Eiselt, A.-K. (2017). Three pillars for the neural control of appetite. Annu. Rev. Physiol. 79, 401–423.

12. Chen, Y., Lin, Y.-C., Kuo, T.-W., and Knight, Z.A. (2015). Sensory detection of food rapidly modulates arcuate feeding circuits. Cell 160, 829–841.

13. Berrios, J., Li, C., Madara, J.C., Garfield, A.S., Steger, J.S., Krashes, M.J., and Lowell, B.B. (2021). Food cue regulation of AGRP hunger neurons guides learning. Nature 595, 695–700.

14. Chen, Y., Lin, Y.-C., Zimmerman, C.A., Essner, R.A., and Knight, Z.A. (2016). Hunger neurons drive feeding through a sustained, positive reinforcement signal. eLife 5.

15. Su, Z., Alhadeff, A.L., and Betley, J.N. (2017). Nutritive, Post-ingestive Signals Are the Primary Regulators of AgRP Neuron Activity. Cell Rep. 21, 2724–2736.

16. Liu, T., Kong, D., Shah, B.P., Ye, C., Koda, S., Saunders, A., Ding, J.B., Yang, Z., Sabatini, B.L., and Lowell, B.B. (2012). Fasting activation of AgRP neurons requires NMDA receptors and involves spinogenesis and increased excitatory tone. Neuron 73, 511–522.

17. Cabral, A., Fernandez, G., Tolosa, M.J., Rey Moggia, Á., Calfa, G., De Francesco, P.N., and Perello, M. (2020). Fasting induces remodeling of the orexigenic projections from the arcuate nucleus to the hypothalamic paraventricular nucleus, in a growth hormone secretagogue receptor-dependent manner. Mol. Metab. 32, 69–84.

18. Wei, W., Pham, K., Gammons, J.W., Sutherland, D., Liu, Y., Smith, A., Kaczorowski, C.C., and O’Connell, K.M.S. (2015). Diet composition, not calorie intake, rapidly alters intrinsic excitability of hypothalamic AgRP/NPY neurons in mice. Sci. Rep. 5, 16810.

19. Park, H., and Poo, M. (2013). Neurotrophin regulation of neural circuit development and function. Nat. Rev. Neurosci. 14, 7–23.

20. Bramham, C.R., and Messaoudi, E. (2005). BDNF function in adult synaptic plasticity: the synaptic consolidation hypothesis. Prog. Neurobiol. 76, 99–125.

21. Unger, T.J., Calderon, G.A., Bradley, L.C., Sena-Esteves, M., and Rios, M. (2007). Selective deletion of Bdnf in the ventromedial and dorsomedial hypothalamus of adult mice results in hyperphagic behavior and obesity. J. Neurosci. 27, 14265–14274.

22. Podyma, B., Johnson, D.-A., Sipe, L., Remcho, T.P., Battin, K., Liu, Y., Yoon, S.O., Deppmann, C.D., and Güler, A.D. (2020). The p75 neurotrophin receptor in AgRP neurons is necessary for homeostatic feeding and food anticipation. eLife 9.

23. Xu, B., Goulding, E.H., Zang, K., Cepoi, D., Cone, R.D., Jones, K.R., Tecott, L.H., and Reichardt, L.F. (2003). Brain-derived neurotrophic factor regulates energy balance downstream of melanocortin-4 receptor. Nat. Neurosci. 6, 736– 742.

24. Molteni, R., Barnard, R.J., Ying, Z., Roberts, C.K., and Gómez-Pinilla, F. (2002). A high-fat, refined sugar diet reduces hippocampal brain-derived neurotrophic factor, neuronal plasticity, and learning. Neuroscience 112, 803–814.

25. An, J.J., Liao, G.-Y., Kinney, C.E., Sahibzada, N., and Xu, B. (2015). Discrete BDNF neurons in the paraventricular hypothalamus control feeding and energy expenditure. Cell Metab. 22, 175–188.

26. Ameroso, D., Meng, A., Chen, S., Felsted, J., Dulla, C.G., and Rios, M. (2022). Astrocytic BDNF signaling within the ventromedial hypothalamus regulates energy homeostasis. Nat. Metab. 4, 627–643.

27. Noble, E.E., Billington, C.J., Kotz, C.M., and Wang, C. (2011). The lighter side of BDNF. Am. J. Physiol. Regul. Integr. Comp. Physiol. 300, R1053–69.

28. Podyma, B., Parekh, K., Güler, A.D., and Deppmann, C.D. (2021). Metabolic homeostasis via BDNF and its receptors. Trends Endocrinol. Metab. 32, 488– 499.

29. Gray, J., Yeo, G.S.H., Cox, J.J., Morton, J., Adlam, A.-L.R., Keogh, J.M., Yanovski, J.A., El Gharbawy, A., Han, J.C., Tung, Y.C.L., et al. (2006). Hyperphagia, severe obesity, impaired cognitive function, and hyperactivity associated with functional loss of one copy of the brain-derived neurotrophic factor (BDNF) gene. Diabetes 55, 3366–3371.

30. Dardennes, R.M., Zizzari, P., Tolle, V., Foulon, C., Kipman, A., Romo, L., Iancu-Gontard, D., Boni, C., Sinet, P.-M., Thérèse Bluet, M., et al. (2007). Family trios analysis of common polymorphisms in the obestatin/ghrelin, BDNF and AGRP genes in patients with Anorexia nervosa: association with subtype, body-mass index, severity and age of onset. Psychoneuroendocrinology 32, 106–113.

31. Reichardt, L.F. (2006). Neurotrophin-regulated signalling pathways. Philos. Trans. R. Soc. Lond. B Biol. Sci. 361, 1545–1564.

32. Underwood, C.K., and Coulson, E.J. (2008). The p75 neurotrophin receptor. Int. J. Biochem. Cell Biol. 40, 1664–1668.

33. Deinhardt, K., and Chao, M.V. (2014). Shaping neurons: Long and short range effects of mature and proBDNF signalling upon neuronal structure. Neuropharmacology 76 Pt C, 603–609.

34. Betley, J.N., Xu, S., Cao, Z.F.H., Gong, R., Magnus, C.J., Yu, Y., and Sternson, S.M. (2015). Neurons for hunger and thirst transmit a negative-valence teaching signal. Nature 521, 180–185.

35. Mazzone, C.M., Liang-Guallpa, J., Li, C., Wolcott, N.S., Boone, M.H., Southern, M., Kobzar, N.P., Salgado, I. de A., Reddy, D.M., Sun, F., et al. (2020). High-fat food biases hypothalamic and mesolimbic expression of consummatory drives. Nat. Neurosci. 23, 1253–1266.

36. Beutler, L.R., Corpuz, T.V., Ahn, J.S., Kosar, S., Song, W., Chen, Y., and Knight, Z.A. (2020). Obesity causes selective and long-lasting desensitization of AgRP neurons to dietary fat. eLife 9.

37. Lu, B. (2003). BDNF and activity-dependent synaptic modulation. Learn. Mem. 10, 86–98.

38. Lohof, A.M., Ip, N.Y., and Poo, M.M. (1993). Potentiation of developing neuromuscular synapses by the neurotrophins NT-3 and BDNF. Nature 363, 350–353.

39. Kang, H., and Schuman, E.M. (1995). Long-lasting neurotrophin-induced enhancement of synaptic transmission in the adult hippocampus. Science 267, 1658–1662.

40. Levine, E.S., Dreyfus, C.F., Black, I.B., and Plummer, M.R. (1995). Brain-derived neurotrophic factor rapidly enhances synaptic transmission in hippocampal neurons via postsynaptic tyrosine kinase receptors. Proc Natl Acad Sci USA 92, 8074–8077.

41. Schuman, E.M. (1999). Neurotrophin regulation of synaptic transmission. Curr. Opin. Neurobiol. 9, 105–109.

42. Schinder, A.F., and Poo, M. (2000). The neurotrophin hypothesis for synaptic plasticity. Trends Neurosci. 23, 639–645.

43. Wu, S.-W., and Xu, B. (2022). Rapid and Lasting Effects of Activating BDNF-Expressing PVH Neurons on Energy Balance. eNeuro 9.

44. Harvey, T., and Rios, M. (2024). The role of BDNF and trkb in the central control of energy and glucose balance: an update. Biomolecules 14.

45. Minichiello, L. (2009). TrkB signalling pathways in LTP and learning. Nat. Rev. Neurosci. 10, 850–860.

46. Yoshii, A., and Constantine-Paton, M. (2010). Postsynaptic BDNF-TrkB signaling in synapse maturation, plasticity, and disease. Dev. Neurobiol. 70, 304–322.

47. Kraemer, B.R., Yoon, S.O., and Carter, B.D. (2014). The biological functions and signaling mechanisms of the p75 neurotrophin receptor. Handb. Exp. Pharmacol. 220, 121–164.

48. Colley, B.S., Biju, K.C., Visegrady, A., Campbell, S., and Fadool, D.A. (2007). Neurotrophin B receptor kinase increases Kv subfamily member 1.3 (Kv1.3) ion channel half-life and surface expression. Neuroscience 144, 531–546.

49. Winkel, F., Ryazantseva, M., Voigt, M.B., Didio, G., Lilja, A., Llach Pou, M., Steinzeig, A., Harkki, J., Englund, J., Khirug, S., et al. (2021). Pharmacological and optical activation of TrkB in Parvalbumin interneurons regulate intrinsic states to orchestrate cortical plasticity. Mol. Psychiatry 26, 7247–7256.

50. Yamashita, T., and Tohyama, M. (2003). The p75 receptor acts as a displacement factor that releases Rho from Rho-GDI. Nat. Neurosci. 6, 461–467.

51. Zhang, Y.H., Chi, X.X., and Nicol, G.D. (2008). Brain-derived neurotrophic factor enhances the excitability of rat sensory neurons through activation of the p75 neurotrophin receptor and the sphingomyelin pathway. J Physiol (Lond) 586, 3113–3127.

52. Sumithran, P., Prendergast, L.A., Delbridge, E., Purcell, K., Shulkes, A., Kriketos, A., and Proietto, J. (2011). Long-term persistence of hormonal adaptations to weight loss. N. Engl. J. Med. 365, 1597–1604.

53. Steuernagel, L., Lam, B.Y.H., Klemm, P., Dowsett, G.K.C., Bauder, C.A., Tadross, J.A., Hitschfeld, T.S., Del Rio Martin, A., Chen, W., de Solis, A.J., et al. (2022). HypoMap-a unified single-cell gene expression atlas of the murine hypothalamus. Nat. Metab. 4, 1402–1419.

